# Influence of Salt Stress on the flg22 Induced ROS Production in *Arabidopsis Thaliana* Leaves

**DOI:** 10.1101/727040

**Authors:** Amrahov Nurlan Rashid, Martin Janda, Mammadov Ziaddin Mahmud, Olga Valentová, Lenka Burketová, Quliyev Akif Alekber

**Affiliations:** Department of Biochemistry and Biotechnology, Baku State University, Baku, Azerbaijan; Institute of Experimental Botany of the Czech Academy of Sciences, Czech Republic; Department of Biochemistry and Microbiology, University of Chemistry and Technology, Prague, Czech Republic

**Keywords:** *Arabidopsis thaliana*, flg22, salt stress, ROS production

## Abstract

In their natural habitats, plants have to cope with multiple stress factors triggering respective response pathways, leading to mutual interference. Our work aimed to study the effect of salt stress in combination with immune response triggered by microbe-associated molecular pattern (MAMP) in *Arabidopsis thaliana* Col-0 plants. We measured ROS production after treatment with flg22 and the influence of concomitant salt stress (NaCl and Na_2_CO_3_).The maximum combined effect of NaCl solution and flg22 on ROS production was achieved at 6 mM salt, which was almost 2 times higher than the single effect of MAMP. A similar maximum combined effect with Na_2_CO_3_ was observed at 10 mM concentration. High concentration of NaCl and Na_2_CO_3_ was accompanied with declining of ROS production, which was completely inhibited at 150 mM of NaCl and at 50 mM of Na_2_CO_3_.The immediate and long term (24 h) effect of NaCl on leaf tissue of *Arabidopsis thaliana* showed that the impact of salt stress on flg22induced ROS production probably did not affect the genetic aspects of the tissue response, but was associated with ionic and osmotic stresses. Experiments with mannitol, KCl and CaCl_2_ allowed to conclude that the observed effect was due to the ionic stress of the salt rather than the osmotic one.

## Introduction

One of the most common and widely distributed factors in the world that negatively impact on the growth and development of plants, including its productivity, is the salinity of the environment (1,2). Since plants have sessile lifestyle, they are not able to escape from the extreme habitat in which they are located, they have to adapt to this environment to survive. Plants have evolved complex protective mechanisms aimed to prevent these combined negative effects and have created opportunities to adapt to these conditions (3, 4). Despite the fact that adaptive responses are complex and multi-stage, they have universal nature and begin with the launch of some biochemical processes. One of them is one of the most important reactions in induction of ROS (reactive oxygen species) production (5), closely related to the process of transmission electrons from NADPH to O_2_whichis catalyzed by the membrane-associated NADPH-oxidase enzyme (6, 7),in plants called respiratory burst oxidase homolog D (RbohD).In most cases, along with abiotic stress, plants are affected by biotic stress. In *Arabidopsis thaliana* plant recognition of pathogen could be realized by its flagellin-sensing 2 (FLS2) receptor kinase which recognizes bacterial flagellin,. After recognition FLS2 binds to co-receptor BAK1 and triggers the events leading to transient high ROS production. Our purpose was to investigate the influence of salt stress (NaCl and Na_2_CO_3_)on the ROS production in *Arabidopsis thaliana* Col-0 leaves in the presence of the elicitor flg22 (conserved N terminal 22 AA polypeptide of flagellin).

## Material and methods

### Plant material

Experiments were carried out on the seedlings of *Arabidopsis thaliana* plants, (ecotype Col-0). Seeds of *A.thaliana* were sterilized by 30% bleach (containing HCl and Tween) and were sown in Jiffy 7 peat pellets. Seedlings were cultivated under a short-day photoperiod (10h/14h light/dark regime)at 100-130 μE m^−2^ s^−1^, 22°C and 70% relative humidity in Snijders plant growth chamber. They were watered with fertilizer-free distilled water as necessary. After 10 days the seedlings were replaced to the new pot with soil and were grown for 4 weeks. For analyses discs 3 mm in diameter, prepared from the leaves of four week old plants were used.

### Chemical treatments

Discs were incubated in 96-well plate in 200 μl of distilled water (control), 6 mM and 150 mM NaCl or 10 mM and 50 mM Na_2_CO_3_ solutions. D-mannitol at concentrations −0,1 mM,1 mM,5mM, 10 mM, 50 mM, 100 mM, 300mM were used as osmotic inducers, where discs were incubated for 20 h. 6 mM, 30mM, 150 mM of KCl and 0,1 mM, 1 mM, 30 mM,150 mM of CaCl_2_ were applied as comparative ion stressor factor. The measurement of hydrogen peroxide production was performed for a period of 50-60 min immediately after adding100 nMflg22 on the discs in all cases (for immediate or 20 hours analysis), replaced in reaction mixture. In order to understand the influence of salt stress on the production of ROS in the presence of flg22, we used different concentrations of stressors in two modifications:

### Hydrogen peroxide determination

H_2_O_2_ production was determined by the luminol-based assay (11,24). This method is based on the detection of the luminescence released by excited luminol molecules generated after the horseradish peroxidase (HRP), which catalyze oxidation of luminol in the presence of H_2_O_2_. Levels as well as duration of the luminescence are proportional to the amount of H_2_O_2_ produced by elicited leaf discs.

For realization the hydrogen peroxide determination discs were replaced in 96-well plate containing 200 μl of reaction mixture, consisting of 17 μg/mL of luminol, 10 μg/mL of horseradish peroxidase (Sigma, P-8125) and 100 nM of flg22.The measurement was performed with a luminometer (Tecan Infinite F200).

## Results

The goal of our study was to determine the impact of salt (NaCl) and alkaline stress (Na_2_CO_3_) on the production of ROS, which is synthesized with the participation of the biotic stress factor. By this way we tried to find out the cause of the induction or repression of ROS products under the influence of these combined stresses with the immediate and delayed action of the abiotic stress factors + biotic stressor.

1. 20 hours analysis. The leaf discs were placed in microplates with solutions containing 6 mM or 150 mM of NaCl, 10 mM or 50 mM of Na_2_CO_3_, 0,1 mM,1 mM, 5mM, 10 mM, 50 mM, 100 mM, 300mMconcentrations of mannitol, 6 mM, 30mM, 150 mM of KCI and 0,1 mM, 1 mM, 30 mM, 150 mM of CaCl_2_ for 20 hours (distilled water was used as a control). Afterwards, the saline and distilled water from the microplate were replaced by a reaction medium to determine the presence of hydrogen peroxide.
2. Immediate analysis. The leaf discs were placed in distilled water for 20 hours. Immediately before determination of hydrogen peroxide .were placed the distilled water and added the solution of 10 mM and 50 mM of NaCl to the reaction mixture.

According to the intensity of the relative luminescence, the use of low concentration of NaCl (6 mM) enhanced the production of ROS compared to controls, both with immediate (fig. 1 A) and after 20 hours of NaCl treatment analysis (fig. 1B). The luminescence intensity in both cases was similar. Soit gradually increased, reached its maximumat10-14 min, then gradually decreased and stopped completely after approximately 30 min. At the maximum values of luminescence under NaCl salt stress, the level of luminescence in immediate experiments increased 2.06 times as compared with the control, but in 20 hours experiments this increase was 1.83 times. Compare with the control (biotic stressor) high concentrations of NaCl (150 mM) totally blocked the synthesis of ROS both after immediate application of NaCl or 20 hours of pre-treatment (fig. 1C, 1D respectively), actually the stimulating effect flg22 under biotic stressor on ROS syntheses disappeared too.

**Fig. 1.**
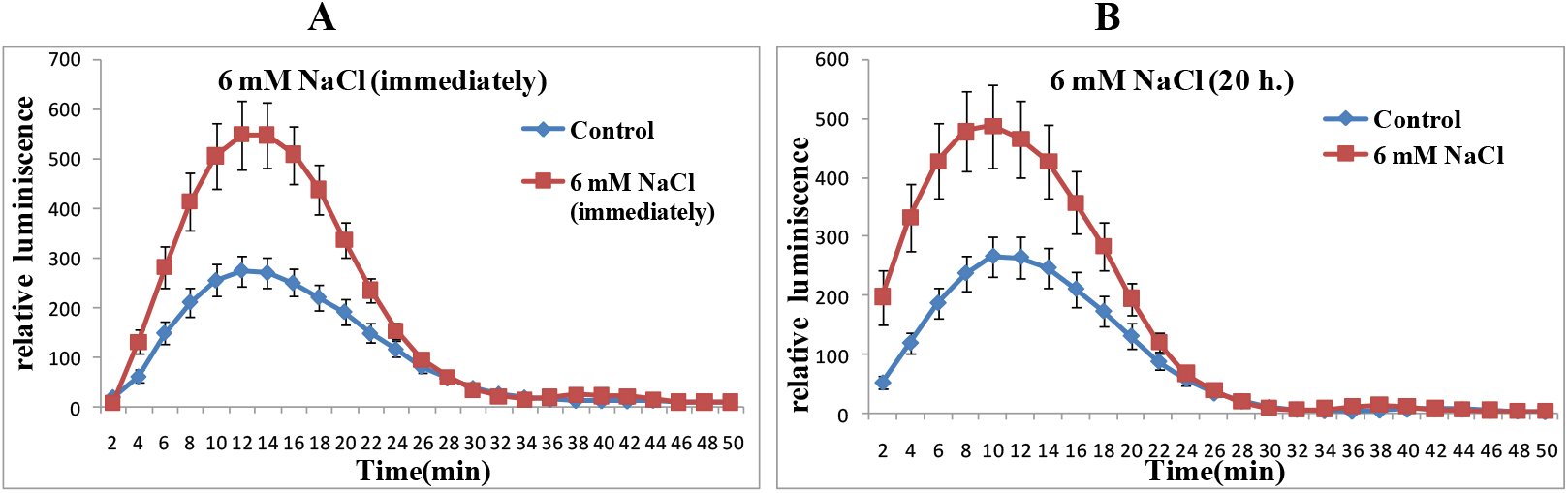
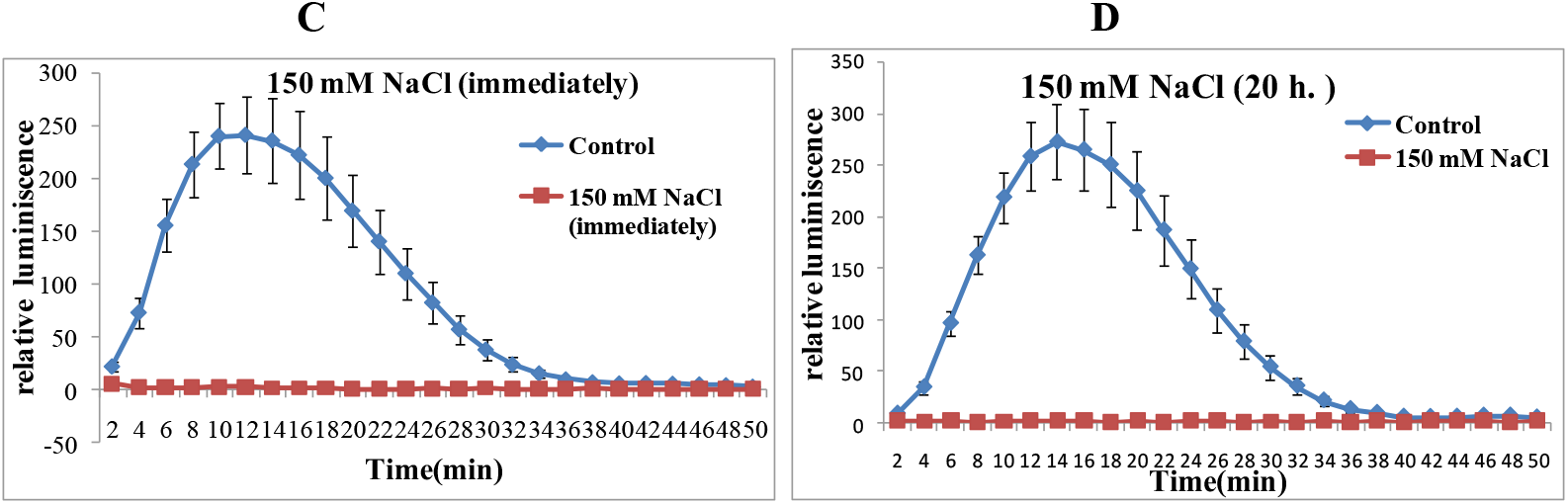
ROS production of *A. thaliana* leaf discs under NaCI stress in the presence of flg22. Data represents mean±SE (standard error) from 12 independent samples. The experiment was done in two biological replicates with similar results. Both types of samples (control and treated ones) were treated with 100 nM flg22.

Low concentration (10mM) of Na_2_CO_3_also enhanced the production of ROS triggered by flg22 (Fig. 2 A) substantially, whereas the high concentration of it (50 mM) blocked the synthesis of ROS completely, suppressing in addition to it the effect of flg22 (Fig. 2 B). Checking the pH of the Na_2_CO_3_solutions showed that the pH of both solutions was only slightly different (pH of 50 mM Na_2_CO_3_ was 11,1 and pH of 10 mM Na_2_CO_3_ was 11,01). According to this fact, we can say that the induction of ROS synthesis was not a consequence of the pH of the medium.

**Fig. 2.**
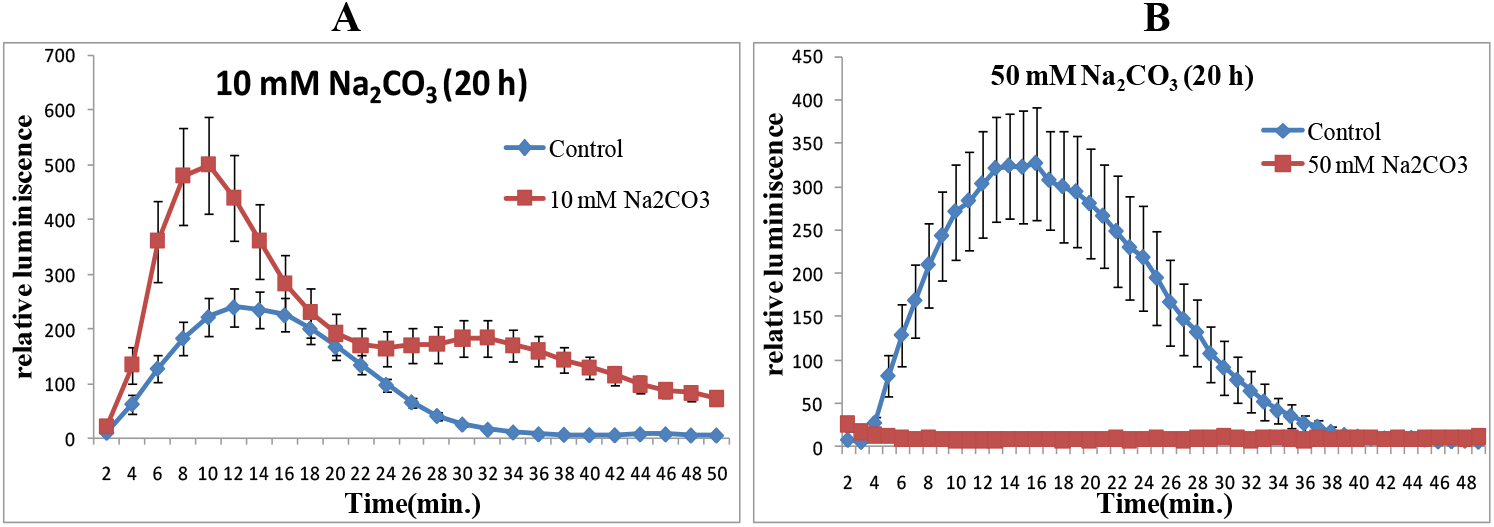
ROS production of *A. thaliana* leaf discs under Na_2_CO_3_stress in the presence of flg22. Data represents mean±SE (standard error) from 12 independent samples. The experiment was done in two biological replicates with similar results. Both types of samples (control and treated ones) were treated with 100 nM flg22.

Salts cause osmotic stress and ion imbalance, which leads to disturbance of membrane ion homeostasis. In order to understand the primary reason of ROS production under the influence of salt stress in the presence of flg22, we tried to investigate this process in more detail. We set a goal to study both of these factors on the ROS production separately. For this purpose, D-mannitol were used as osmotic stress factor, whereas KCl and CaCl_2_as ionic stressors.

The results of the study on the effect of various concentrations of mannitol on the ROS production are presented in Fig. 3. Osmotic stress, caused by mannitol within the concentration 0.1-300 mM did not induce the ROS production, on the contrary, depending on the concentration the synthesis was inhibited directly (Fig.3 A and 3B).

**Fig. 3.**
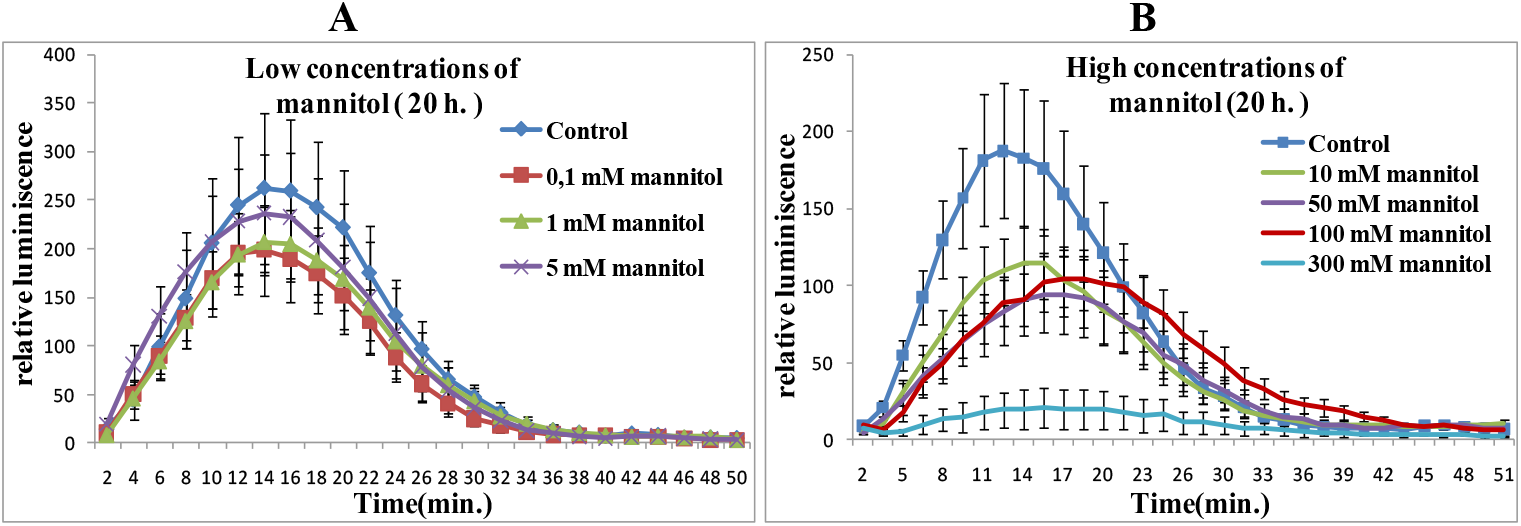
ROS production of *Arabidopsis thaliana* leaf discs under osmotic stress in the presence of flg22. Data represents mean±SE (standard error) from 12 independent samples. The experiment was done in two biological replicates with similar results. Both types of samples (control and treated ones) were treated with 100 nM flg22.

Low concentrations of KCl led to the increase of ROS production, while high concentrations of the salt had an inhibitory effect (Fig. 4 A). This made it clear that with the participation of flg22, Na-K homeostasis was one of the driving forces in the activation and inhibition of ROS synthesis. In comparative experiments with identical concentrations of NaCl and KCl, it was found that both ions exhibited the same effect, contributing to an increase in the synthesis of ROS (with the participation of flg22), compared to the control (only flg22).This findings again proved our assumption of Na-K homeostasis and its influence on the production of ROS after flg22 treatment (Fig. 4B). It was also demonstrated that CaCl_2_ had a similar effect as NaCl and KCl. Low concentration ofCaCI_2_led to the increase, while higher concentrations caused the decrease of ROS production. At 150 mM of CaCI_2_ ROS synthesis was blocked totally (Fig. 4 C, 4 D).

**Fig. 4.**
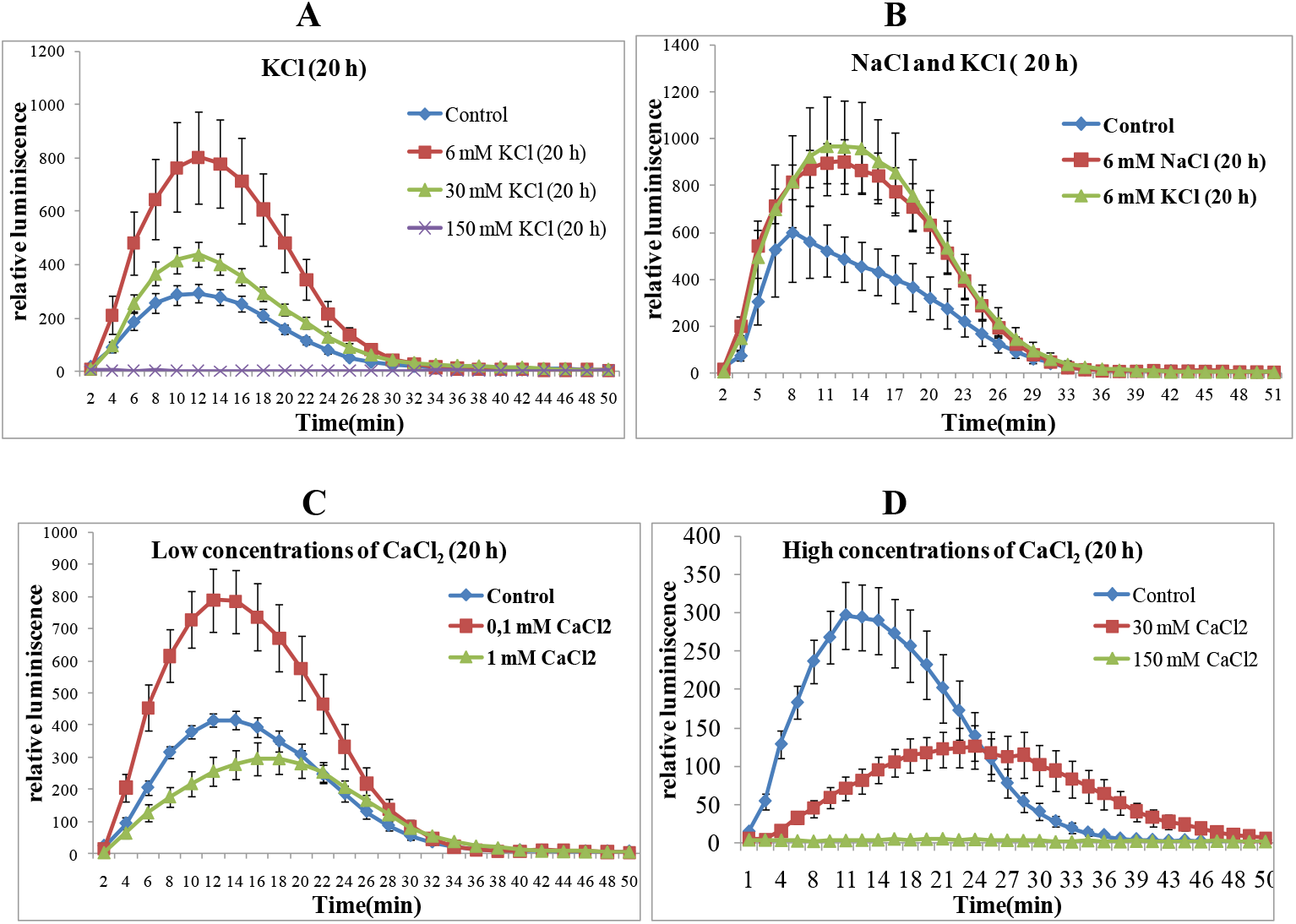
ROS production of *Arabidopsis thaliana* leaf discs under ionic stress in the presence of flg22. Data represents mean±SE (standard error) from 12 independent samples. The experiment was done in two biological replicates with similar results. Both types of samples (control and treated ones) were treated with 100 nM flg22.

## Discussion

In our study, non-genetic mechanisms of activation of ROS were investigated.

We have determined that low concentrations of salt (6 mM NaCl) and alkaline stress (10 mM Na_2_CO_3_) in the presence of biotic stress (flg22) lead to increased ROS production. At the same time, under similar biotic stress factor, high salinization (150 mM NaCl) and alkalization (50 mM Na_2_CO_3_) were accompanied with a decrease in the production of ROS.

Salt stress, i.e. abiotic stress leads to a disturbance of the Na / K-homeostasis. Ionic stress caused by partial accumulation of Na^+^ in the extracellular space leads to partial inhibition of Na^+^/K^+^ ATP–ase. As a result, the amount of Na^+^ in the intracellular space increases. And to prevent this phenomenon, in a very short time, the concentration of Ca^2+^ in the intracellular space increases, which significantly reduces Na^+^ toxicity (16, 17).

The effect of biotic stress could be explained as follows.

It is known that the flagellin-sensing2 (FLS2) receptor kinase, which belong to leucine-rich repeat receptor kinases (LRR-RKs), recognizes flg22, that is 22 AAN-terminal peptide of conserved bacterial protein-flagellin. Subsequently, the intracellular domain-BAK1, which is responsible for the transmission of the activation signal to RbohD, was involved in enzyme catalyzing and triggering ROS production(8–10). Activation of BAK1 leads to increased Ca^2+^ influx by activating GLR3.3. and GLR3.6. channels. An increase in the concentration of intracellular Ca^2+^ affects the TPC1 channel located on the surface of the vacuoles and in turn increases the concentration of Ca^2+^ which begins to flow additionally from the vacuole into the cytoplasm (18).

The immediate effect of salt stresses + biotic stressor that leading to change of ROS production gives us reason to suppose that the process involves clathrin-mediated endocytosis (CME) (12) and Na-K homeostasis.

As it is already known, salt stress, in particular, sodium-dependent salt stress causes various effects, such as ionic toxicity, oxidative and osmotic stresses (13). The increase in extracellular exogenous sodium ions leads to the disruption of sodium-potassium homeostasis and ROS synthesis occurs. In the moderate stress conditions ROS production and potassium leakage could play an essential role as a ‘metabolic switch’ in anabolic reactions, stimulating catabolic processes and saving ‘metabolic’ energy for adaptation and repair needs. But in strong stress conditions these processes could lead to programmed cell death (14,15).

Thus, RbohD under low concentrations of salt, i.e. 6 mM NaCl, 6 mM KCl, 0,1mM CaCl_2_ and 10 mM Na_2_CO_3_ were subjected to two influences. First one is the effect of the signal that comes from the interaction of flg22-FLS2-BAK1-BIK1-RbohD, whereas second one is the involving of additional signal from salt-RbohD interaction. Since, at low concentrations of salt (6 mM NaCl) and alkaline stress (10 mM Na_2_CO_3_,) a change in the number of ROS was not observed, and according to literature sources, there was also an increase in the concentration of intracellular Ca^2+^, we assume that biotic stress plays a key role in starting ROS production. Under the influence of biotic stress, along with an increase in Ca^2+^ concentration inside the cell, a cascade mechanism is activated to activate the key domains of BAK1, and then BIK1, associated with RbohD. Thus, the activation of the domain BIK1 and launches products ROS. (19, 20).

Ca^2+^, by binding with the EF-hand of NADPH-oxidase, leads to the production of ROS (21–23).

As already described by Hao H. et al (12), treatment of plants with 100 mM of NaCl (high concentration) resulted in an increased endocytosis of RbohD and reducing of RbohD on the membrane surface. At the same time a high concentration of salt stress led to degradation of RbohD. In our experiment, high concentrations of salts (150 mM NaCl, 50 mM Na_2_CO_3_, 150 mM KCl and 150 mM CaCl_2_) also led to a decrease in ROS production, which gave us an understanding of the involvement of endocytosis and degradation as negative factors in ROS production.

High salt concentration also led to an osmotic stress. Using D-mannitol as an osmotic stressor, we observed a decrease in ROS production. Consequently, a strong osmotic stress caused by high concentrations of salts or mannitol, combined with biotic stress, leads to a decrease in the production of ROS, and therefore also a weakening of the protective reaction.

## Conclusion

Salt stress, created by low concentrations of NaCl and Na_2_CO_3_in the presence of the elicitor flg22 induced the ROS production in *Arabidopsis thaliana* leaf tissues. Induced ROS production probably affected as the genetic transcriptomic changes aspects of in the tissue response, but also associated with ionic stress. This effect was caused by the changes in ionic strength rather than the osmotic effect.

## Acknowledgments

The work was supported from the International Visegrad Fund, Visegrad Scholarship Program / V4EaP Scholarship Program, under the grant N° 51701351.

## Notes

#### Summary of Updates

Our work aimed to study the effect of salt stress in combination with immune response triggered by microbe-associated molecular pattern (MAMP) in Arabidopsis thaliana Col-0 plants. We measured ROS production after treatment with flg22 and the influence of concomitant salt stress (NaCl and Na2CO3).The maximum combined effect of NaCl solution and flg22 on ROS production was achieved at 6 mM salt, which was almost 2 times higher than the single effect of MAMP. A similar maximum combined effect with Na2CO3 was observed at 10 mM concentration. High concentration of NaCl and Na2CO3 was accompanied with declining of ROS production, which was completely inhibited at 150 mM of NaCl and at 50 mM of Na2CO3.The immediate and long term (24 h) effect of NaCl on leaf tissue of Arabidopsis thaliana showed that the impact of salt stress on flg22 induced ROS production probably did not affect the genetic aspects of the tissue response, but was associated with ionic and osmotic stresses. Experiments with mannitol, KCl and CaCl2 allowed to conclude that the observed effect was due to the ionic stress of the salt rather than the osmotic one.

